# A genome-scale Opisthokonta tree of life: toward phylogenomic resolution of ancient divergences

**DOI:** 10.1101/2023.09.20.556338

**Authors:** Hongyue Liu, Jacob L. Steenwyk, Xiaofan Zhou, Darrin T. Schultz, Kevin M. Kocot, Xing-Xing Shen, Antonis Rokas, Yuanning Li

## Abstract

Ancient divergences within Opisthokonta—a major lineage that includes organisms in the kingdoms Animalia, Fungi, and their unicellular relatives— remain contentious, hindering investigations of the evolutionary processes that gave rise to two kingdoms and the repeated emergence of iconic phenotypes like multicellularity. Here, we use genome-scale amounts of data to reconstruct the most taxon-rich Opisthokonta tree of life to date (348 species) and place divergences in geologic time, suggesting a Mesoproterozoic origin (∼ 1.11 billion years ago). By dissecting multiple dimensions of phylogenomic error, such as the influence of taxon sampling and model complexity, we found that deep divergences within Holozoa remain unresolved and suggest Pluriformea is either sister to Ichthyosporea and Filozoa (Pluriformea-sister hypothesis) or is monophyletic to Ichthyosporea, forming the Teretosporea lineage (Teretosporea-sister hypothesis). A combination of information theory and sensitivity analyses revealed that the inferred unicellular Holozoa relationships are largely robust to common sources of analytical error, such as insufficient model complexity, and suggest that previous reports likely suffered from insufficient taxon sampling. Our study presents a robust Opisthokonta phylogenomic framework, highlights the challenges in resolving the relationships of unicellular Holozoa, and paves the way for illuminating ancient evolutionary episodes concerning the origin of two kingdoms.

## Introduction

Eukaryotic multicellularity has evolved independently in several key lineages, including within Opisthokonta, a monophyletic supergroup containing animals, fungi, and their unicellular relatives (Fig. 1)^1–3^. Opisthokonta is divided into two main lineages: Holomycota^4^, containing Fungi and their unicellular relatives (e.g. Nucleariida), and Holozoa^5,6^, which includes Metazoa (Porifera, Placozoa, Ctenophora, Cnidaria, Bilateria) and their unicellular relatives (e.g. Choanoflagellata^7^, Filasterea^8^, Ichthyosporea^9,10^ and Pluriformea^11^; Fig. 2A). Establishing evolutionary relationships among major lineages of Opisthokonta is key for illuminating the origins of animals and fungi, as well as of complex phenotypes like multicellularity^11–18^.

**Fig. 1:**
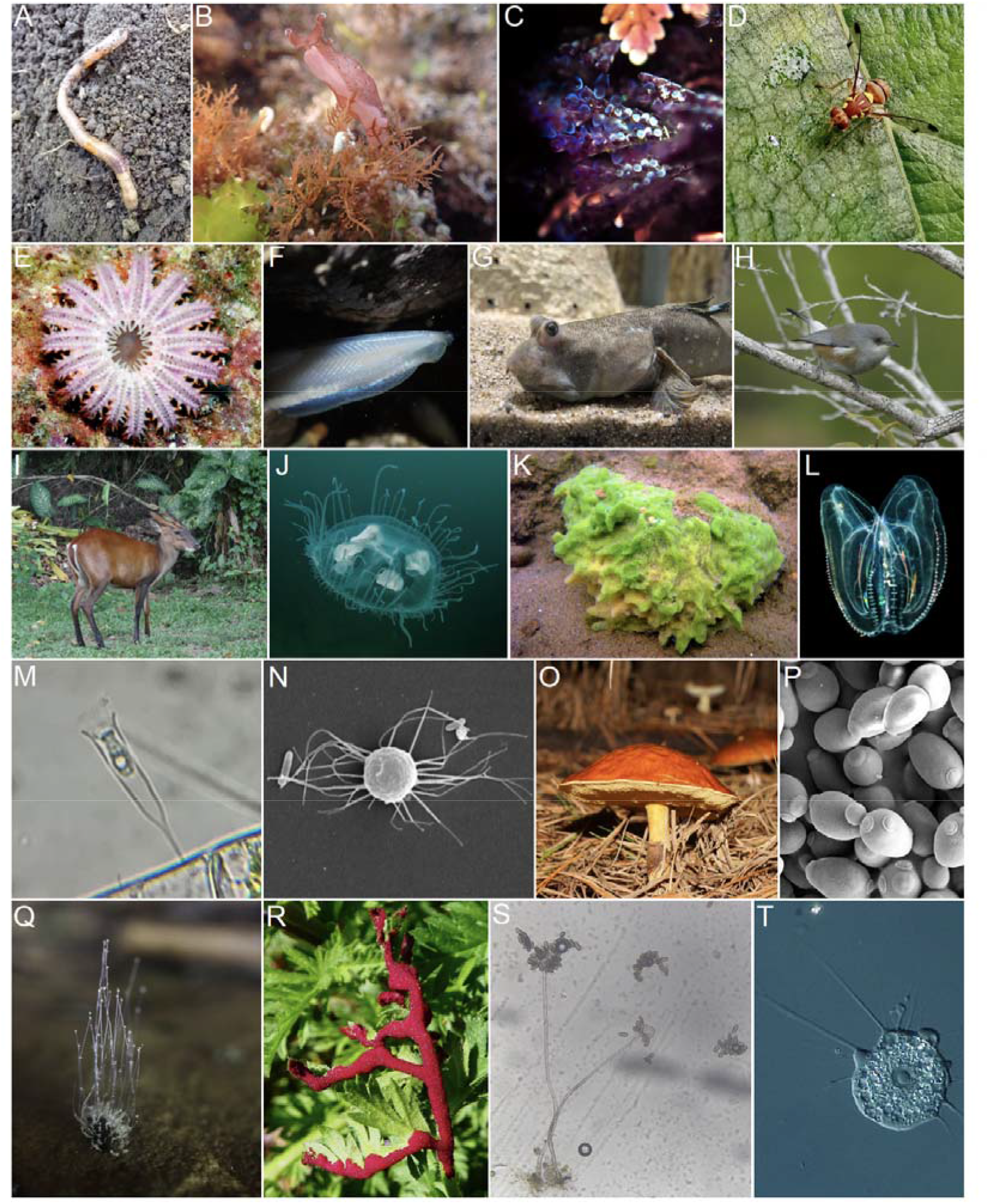
Diversity of major Opisthokonta lineages. (A) Common earthworm *Lumbricus terrestris* (Annelida). (B) California sea hare *Aplysia californica* (Mollusca). (C) Common bugula *Bugula neritina* (Bryozoa). (D) Melon fly *Zeugodacus cucurbitae* (Arthopoda). (E) Crown-of-thorns starfish *Acanthaster planci* (Echinodermata). (F) Lancelets *Epigonichthys hectori* (Cephalochordata). (G) Great blue spotted mudskipper *Boleophthalmus pectinirostris* (Actinopterygii and Chordata). (H) Réunion grey white-eye *Zosterops borbonicus* (Aves and Chordata). (I) Southern red muntjac *Muntiacus muntjak* (Mammalia and Chordata). (J) Peach blossom jellyfish *Craspedacusta sowerbii* (Cnidaria). (K) *Spongilla lacustris* (Porifera). (L) Warty comb jelly *Mnemiopsis leidyi* (Ctenophora). (M) *Salpingoeca gracilis* (Choanoflagellatea). (N) *Ministeria vibrans* (Filasterea). (O) *Suillus luteus* (Basidiomycota). (P) Baker’s yeast *Saccharomyces cerevisiae* (Ascomycota). (Q) *Phycomyces blakesleeanus* (Mucoromycota). (R) *Synchytrium papillatum* (Chytridiomycota). (S) *Rhopalomyces elegans* (Zoopagomycota). (T) *Nuclearia thermophila* (Nucleariida). Images G, N, P, and T are available in the public domain and were sourced from Wikimedia Commons (https://commons.wikimedia.org/wiki/Main_Page). The rest of the images were retrieved from iNaturalist (https://www.inaturalist.org/). All images are credited to various artists under Creative Commons licenses with slight modifications. For specific author names, hyperlinks to the images, and copyright license details, please refer to Supplementary Table 14.

**Fig. 2:**
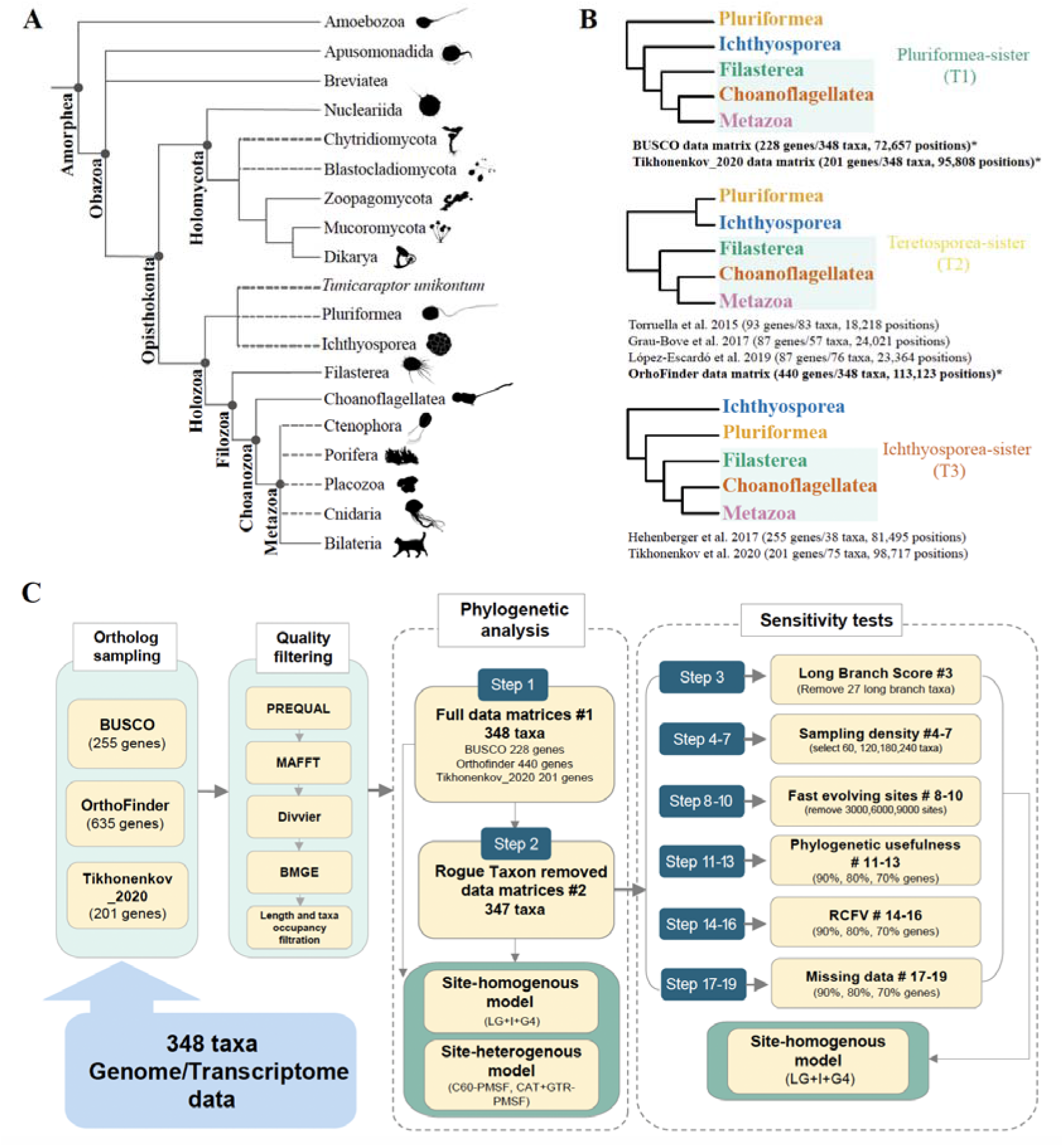
Incongruence across the Opisthokonta Tree of Life and a workflow for examining evolutionary relationships. **A**, Schematic representation of the phylogenetic relationships of Opisthokonta based on recent molecular phylogenies^11,38,39^. Dashed branches reflect uncertain relationships across Opisthokonta. **B**, Alternative hypotheses of the relationships of unicellular holozoans. Studies that support these hypotheses are listed below each tree; studies with an asterisk are results from this study. The three hypotheses from top to bottom are Pluriformea as the sister lineage to the rest of the Holozoa, a clade of Pluriformea + Ichthyosporea as the sister lineage to the rest of the Holozoa, and Ichthyosporea as the sister lineage to the rest of the Holozoa, respectively. **C,** To shed light on these uncertainties, we developed a workflow that broadly samples gene and model space and implements sensitivity analyses to dissect sources of error. Data matrices are referenced throughout the text as BUSCO, OrthoFinder, and Tikhonenkov_2020. Subsampled data matrices have numbers following the ‘#’ character reflecting the filtering step used to generate them. Detailed information on each data matrix can be found in Supplementary Table 2.

In the phylogenomic era, studies utilizing hundreds to thousands of loci have greatly improved our understanding of the Opisthokonta tree of life; however, evolutionary relationships among several key lineages remain disputed (Fig. 2A). Among Holozoa, the root position of the animal tree has emerged as one of the most challenging problems in animal phylogenetics, with either Ctenophora or Porifera placed as the sister lineage to the rest of animals^19–26^. Major clades within Bilateria are also contentious, such as the interrelationships of major subclades within Lophotrochozoa and Ecdysozoa^27–29^. Ambiguity also exists for the phylogenetic placement of Xenacoelomorpha, a clade that contains *Xenoturbella* and Acoelomorpha, as either the sister lineage to the rest of the bilaterians^30–33^ or within Deuterostomia as sister to Ambulacraria^34–36^.

Ancient branching patterns among Holozoa have also proven recalcitrant, wherein different phylogenomic studies support conflicting topologies or are equivocal in support (Fig. 2B). Although previous phylogenomic studies have established a generally stable phylogenetic backbone for the unicellular relatives of animals, supporting Ichthyosporea as the sister lineage to the rest of the Holozoa^8,14,37^ and placing Filasterea in a clade with Choanoflagellata and Metazoa, termed Filozoa^8,14^ (Fig. 2A). However, the recent discovery of novel Holozoa lineages presents new challenges and opportunities for phylogenetic reconstruction. For example, the placement of the newly proposed unicellular lineage Pluriformea, comprised of the enigmatic marine protist *Corallochytrium limacisporum* and the freshwater protist *Syssomonas multiformis*^11^, is contentious; some analyses place it as sister to the Filozoa^11,18^ (Ichthyosporea- sister hypothesis, Fig. 2B), whereas others support a sister relationship to Ichthyosporea, forming the Teretosporea group^38,39^ (Teretosporea-sister hypothesis, Fig. 2B). Similarly, the newly discovered marine protist *Tunicaraptor unikontum* may be sister to Filasterea, Filozoa, or the sister group of the rest of the Holozoa based on phylogenomic analyses^18^.

Among Holomycota, the branching order between flagellated zoospore-producing fungi Blastocladiomycota and Chytridiomycota relative to other fungi remains contentious^26,38,40–43^. Similarly, the phylogenetic placement of the endoparasitic zoosporic fungus *Olpidium* is under debate; *Olpidium* is often placed outside of the core chytrids, closer to terrestrial fungi, but the precise relationships are uncertain^44,45^. These examples illustrate that incongruence is widespread among Opisthokonta, hindering evolutionary studies.

Insufficient taxon sampling is a well-established driver of phylogenomic incongruence and poor support^46,47^. Among shallower, yet still ancient, lineages within Opisthokonta like the Ascomycota, insects, and fish, expanded taxon sampling using hundreds of taxa has improved the resolution of relationships^48–51^. In contrast, phylogenomic datasets so far used to reconstruct the Opisthokonta tree of life tend to have relatively sparse taxon sampling, averaging 63.5 taxa (range: 38 – 83)^11,14,18,38^, raising the hypothesis that improved taxon sampling may ameliorate uncertainty in the Opisthokonta phylogeny.

The flourishing of genomic data offers an opportunity to reconstruct a genome-scale Opisthokonta tree of life, examine its support for relationships that have previously remained poorly resolved, and elucidate drivers of incongruence. Here, we leverage the abundant genomic data across Opisthokonta (348 taxa spanning 33 major lineages, Supplementary Table 1) and reconstruct the first taxon-rich phylogeny of Opisthokonta, with particular emphasis on resolving deep relationships within Holozoa and establishing the timeline of Opisthokonta diversification.

## Results and Discussion

### Phylogenomics uncovers a broadly supported Opisthokonta Tree of Life

To infer the Opisthokonta tree of life, three data matrices (termed BUSCO, OrthoFinder, and Tikhonenkovz_2020, reflecting the origin of phylogenomic markers) with high taxon sampling and gene occupancy were constructed using different orthology inference methods and rigorous quality control measures (Fig. 2C, Table 1, Supplementary Table 2). Summary statistics differ between the data matrices. Approximately half of the genes in each data matrix are not present in the other two data matrices (Extended data Fig. 1); the distribution of functional categories also varies significantly across the matrices, suggesting substantial differences in the biological processes captured by each matrix (Extended data Fig. 1).

**Table 1.** Summary statistics of three phylogenomic data matrices.

| Data matrix | Number of Genes | Number of Sites | Average Taxon Occupancy | Average Gene Length | Average Site Occupancy |
| --- | --- | --- | --- | --- | --- |
| BUSCO | 228 | 72,657 | 79.9% | 319 | 67.1% |
| OrthoFinder | 440 | 113,123 | 85.6% | 257 | 68.9% |
| Tikhonenkov<br>_2020 | 201 | 95,808 | 87.0% | 477 | 72.6% |

The evolutionary history of Opisthokonta was inferred using substitution models with varying complexity. These analyses produced 18 phylogenomic trees: three data matrices (BUSCO, OrthoFinder and Tikhonenkov_2020) * two versions (full data matrix and rogue taxon pruned) * three modeling schemes (LG, LG+C60, GTR+CAT). We found that ∼85% of internal branches were congruent across the 18 trees, suggesting a large fraction of bipartitions in the Opisthokonta phylogeny were robustly supported (Fig. 3A, Supplementary Table 3, Supplementary Figs. 1-18). Within Holozoa, our results recapitulate many deep relationships seen in molecular studies to date: Bilateria, Deuterostomia, Ecdysozoa, Lophotrochozoa, Protostomia and Nephrozoa are all recovered^27–29,52–54^. Our results also consistently support the sister relationship of Filasterea to a Choanoflagellatea and Metazoa group (Filozoa hypothesis^4,8,37^, although this result is not always robustly supported, Fig. 3A, Supplementary Figs. 1-18).

**Fig. 3:**
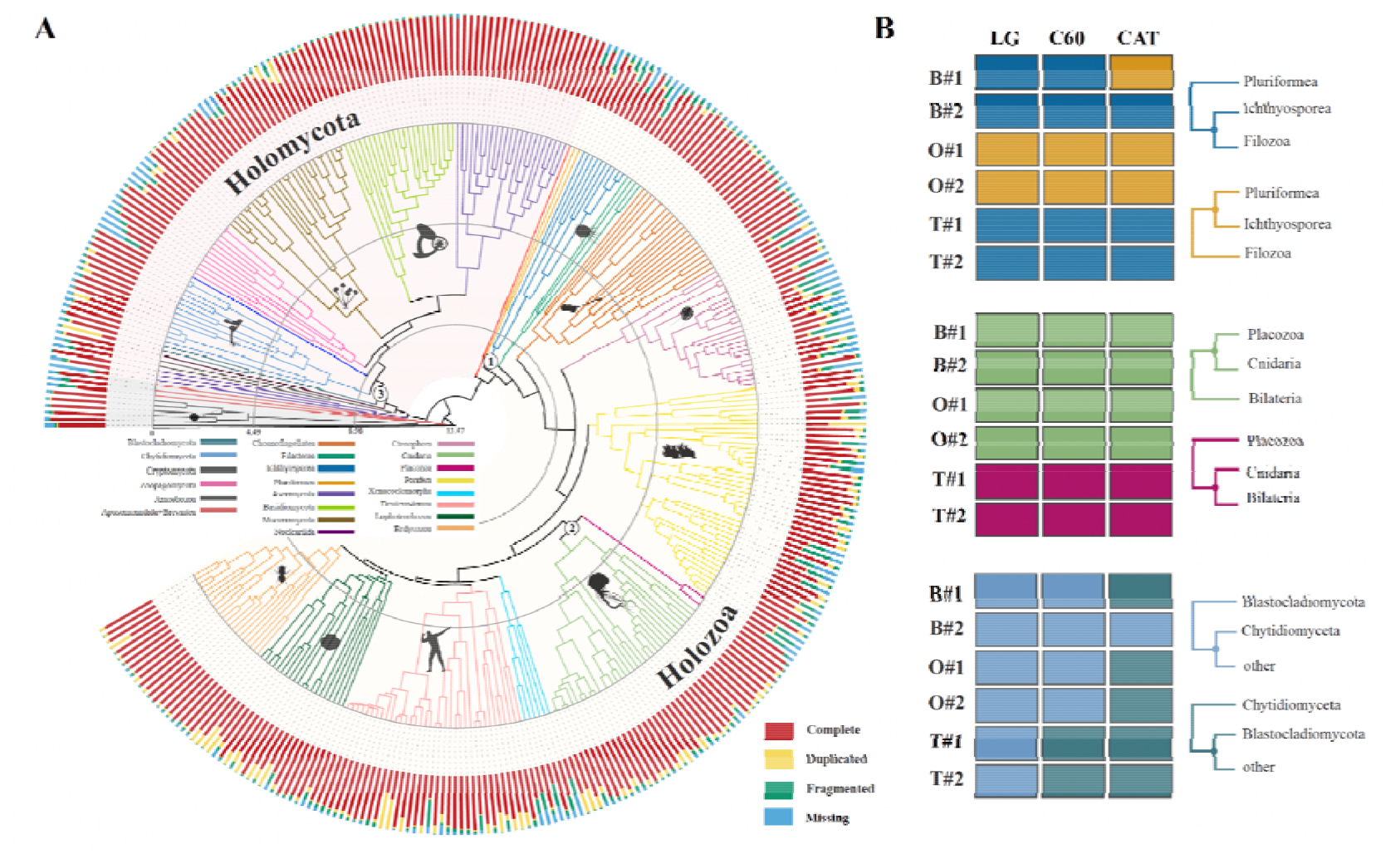
Time-calibrated phylogeny of 348 species spanning the diversity of opisthokonts. **A**, Divergence time estimation using MCMCTree with a topology reconstructed from the concatenation-based maximum likelihood analysis of OrthoFinder#1 data matrix using the LG+C60 model. The bar plot next to each species indicates genomic quality assessed using BUSCO. ‘‘Complete’’ indicates the fraction of full-length BUSCO genes; “Duplicated’’ indicated if there were two or more complete predicted genes for one BUSCO gene, ‘‘Fragmented’’ indicates the fraction of genes with a partial sequence; and ‘‘Missing’’ indicates the fraction of genes not found in the genome. Images from phylopic.org. The timescale is in 100 millions of years before present. See also Extended data Fig. 2, Supplementary Figs. 1-18, 82. **B**, The distribution of topology supported across matrices and evolutionary models, colored according to topology supported. The grids correspond to three contentious nodes indicated in A. “B”, “O”, “T” represents BUSCO, OrthoFinder, and Tikhonenkov_2020, respectively.

Among Holomycota, notable examples of relationships recovered consistently in our results include the monophyly of the Dikarya subkingdom^55^, comprising the Ascomycota and Basidiomycota phyla, which received maximal support. Mucoromycota was recovered as the sister group of Dikarya^42^ and Zoopagomycota is sister to both lineages^56^. Supporting a recent study, a Nucleariida clade consisting of *Parvularia atlantis*, *Fonticula alba*, and *Lithocolla globosa* was recovered as the sister lineage to the rest of the Holomycota^57^ (Fig. 3A, Supplementary Figs. 1-18).

### Opisthokonta diversified in the Mesoproterozoic

A Bayesian relaxed molecular clock calibrated with ten widely accepted fossil calibration points (Supplementary Table S4) facilitated estimating divergence times of Opisthokonta evolution (Fig. 3A, Table 2, Supplementary Fig. 82). These analyses infer that animals and fungi diverged approximately 1112 million years ago (Mya) (95% credibility interval (CI) ranging from 1003 to 1220 Mya). The origin of Holomycota is estimated to be approximately 1030 Mya (95% CI: 921 to 1139 Mya) and Holozoa emerged roughly 1023 Mya (95% CI; 927 to 1120 Mya). The divergence between Choanoflagellata and Metazoa, was estimated to have transpired in the Late Proterozoic Tonian period 889 Mya (95% CI: 812 and 967 Mya). The origin of animals, marking the emergence of animal multicellularity, began approximately 799 Mya (95% CI, 758 to 840 Mya). The estimated divergence time between protostomes and deuterostomes was approximately 613 to 641 Mya (mean: 627). Within Holomycota, the origin of the kingdom Fungi (the sister clade to Nucleariida) was dated to 961 Mya (95% CI, 855 to 1067 Mya), consistent with a previous report using fewer loci^58^. The origin of terrestrial fungi was estimated at 775 Mya (95% CI: 688 to 862 Mya) and the origin of Dikarya was estimated to have occurred around 665.5 Mya (95% CI: 580 to 751 Mya).

**Table 2.** Inferred time intervals for the various Opisthokonta clades, in millions of years before present.

| **Crown group** | **min** | **max** | **width** | **mean** |
| --- | --- | --- | --- | --- |
| Opisthokonta | 1003.2 | 1220.1 | 216.8 | 1111.6 |
| Holomycota | 921 | 1139.2 | 218.2 | 1030.1 |
| Holozoa | 926.8 | 1119.6 | 192.8 | 1023.2 |
| Choanozoa | 811.5 | 967.1 | 155.6 | 889.3 |
| Metazoa | 742.9 | 836.5 | 93.6 | 789.7 |
| Porifera-ParaHoxozoa | 695.6 | 782 | 86.5 | 738.8 |
| Bilateria | 629.4 | 655.6 | 26.2 | 642.5 |
| Deuterostomia | 570.7 | 618.3 | 47.6 | 594.5 |
| Ecdysozoa | 545.4 | 591.7 | 46.3 | 568.5 |
| Lophotrochozoa | 571.6 | 601.9 | 30.4 | 586.8 |
| Chordata | 525.4 | 590 | 64.6 | 557.7 |
| Ecdysozoa-Lophotrochozoa | 593.3 | 623.6 | 30.4 | 608.4 |
| Ascomycota | 443.2 | 635.2 | 192 | 539.2 |
| Basidiomycota | 435.2 | 632.8 | 197.6 | 534 |
| Mucoromycota | 503.1 | 716.2 | 213.1 | 609.65 |
| Zoopagomycota | 633.9 | 815.4 | 181.5 | 724.65 |
| Chytridiomycota | 604.2 | 816.9 | 212.7 | 710.55 |
| Obazoa | 1102.3 | 1360.4 | 258.1 | 1231.3 |

This study involves the first comprehensive phylogenomic timetree of Opisthokonta, filling a gap in the literature. These results may inform the testing of hypotheses that tie the emergence of lineages and phenotypes to specific geologic events. For example, molecular dating analyses have consistently placed the emergence of animals in the Tonian-Cryogenian period, approximately 850 to 635 Mya^59–61^, broadly coinciding with the rise in atmospheric oxygen levels and changes in the phosphorus cycle^62,63^.

### Widely sampling parameter space brings uncertainty into focus

Despite assembling genome-scale data matrices comprising tens of thousands of well-aligned sites from hundreds of taxa and using advanced site-heterogeneous models, we encountered persistent conflicts in phylogenetic relationships arising from different gene sets. Examination of incongruence across the 18 inferred phylogenies revealed that approximately 15% of bipartitions were unstable, particularly affecting historically contentious branches (Fig. 3B, Supplementary Table 3, Supplementary Figs. 1-18).

For instance, one of the conflicting topologies concerned the position of Placozoa (Fig. 3B): Tikhonenkov_2020 matrices support the sister relationship between Cnidaria and Bilateria with Placozoa as sister to this clade. In contrast, the BUSCO and OrthoFinder matrices recovered a sister taxon relationship between Placozoa and Cnidaria; both topologies were reported before^64,65^.

Furthermore, the branching order of Chytridiomycota and Blastocladiomycota also remains elusive and is sensitive to model complexity (Fig. 3B, Supplementary Figs. 1-18); support for Chytridiomycota sister to Blastocladiomycota and other fungi was only observed using complex mixture models. In addition, the placement of *Olpidium* was unstable and data matrix dependent. Consistent with a previous report^45^, the OrthoFinder and Tikhonenkov_2020 data matrices strongly supported *Olpidium* as sister to a clade of non-flagellated terrestrial fungi (Supplementary Figs. 7-18), supporting a single loss event of the fungal flagellum^66^. However, the BUSCO data matrices supported *Olpidium* nested within non-flagellated fungi (Supplementary Figs. 1- 6), either as the sister group of Mucoromycota, or as the sister group to Dikarya, supporting multiple losses of the flagellum.

These findings highlight the complexity of resolving certain relationships in the Opisthokonta tree of life. Undue confidence in tree topology can negatively impact downstream analyses relying on phylogeny, such as studies of phenotype evolution and biomolecules. By widely sampling parameter space (genes and model complexity), uncertainty can be more readily detected^49,50^. Our rigorous approach is broadly applicable to other tree of life inquiries and illuminates evolutionary relationships warranting further investigation.

### Phylogenomics supports novel relationships among unicellular Holozoa

Resolving relationships among major groups of the unicellular Holozoa has proven challenging, with previous studies yielding contrasting topologies or weakly supporting critical nodes, especially regarding the relationships between Ichthyosporea and Pluriformea^11,16,18,39,67^. The present analyses support a novel resolution of unicellular Holozoa suggesting Pluriformea is the sister group to both Ichthyosporea and Filozoa when using the BUSCO and Tikhonenkov_2020 data matrices (Pluriformea-sister hypothesis, Fig. 2B, Extended data Fig. 2A, Supplementary data Figs. 1-6, 13-18). In contrast, the OrthoFinder data matrix suggests that Pluriformea is the sister taxon to Ichthyosporea (known as the Teretosporea group), as reported in previous studies^38,39,67^ (Teretosporea-sister hypotheses, Fig. 2B, Extended data Fig. 2B, Supplementary data Figs. 7-12).

Relationships among unicellular Holozoa are robust to substitution model complexity, except for one instance in which the BUSCO#1 matrix with GTR+CAT model weakly supported T2 (UFP = 23) (Supplementary Fig. 3). The third alternative topology, which supports Ichthyosporea as the sister taxon to all other Holozoa (Ichthyosporea-sister hypothesis, Fig. 2B)^11,18^ was not recovered in our analyses and was rarely observed among UFB approximated trees, suggestive of minimal support (Fig. 4A, Supplementary Table 5).

**Fig. 4:**
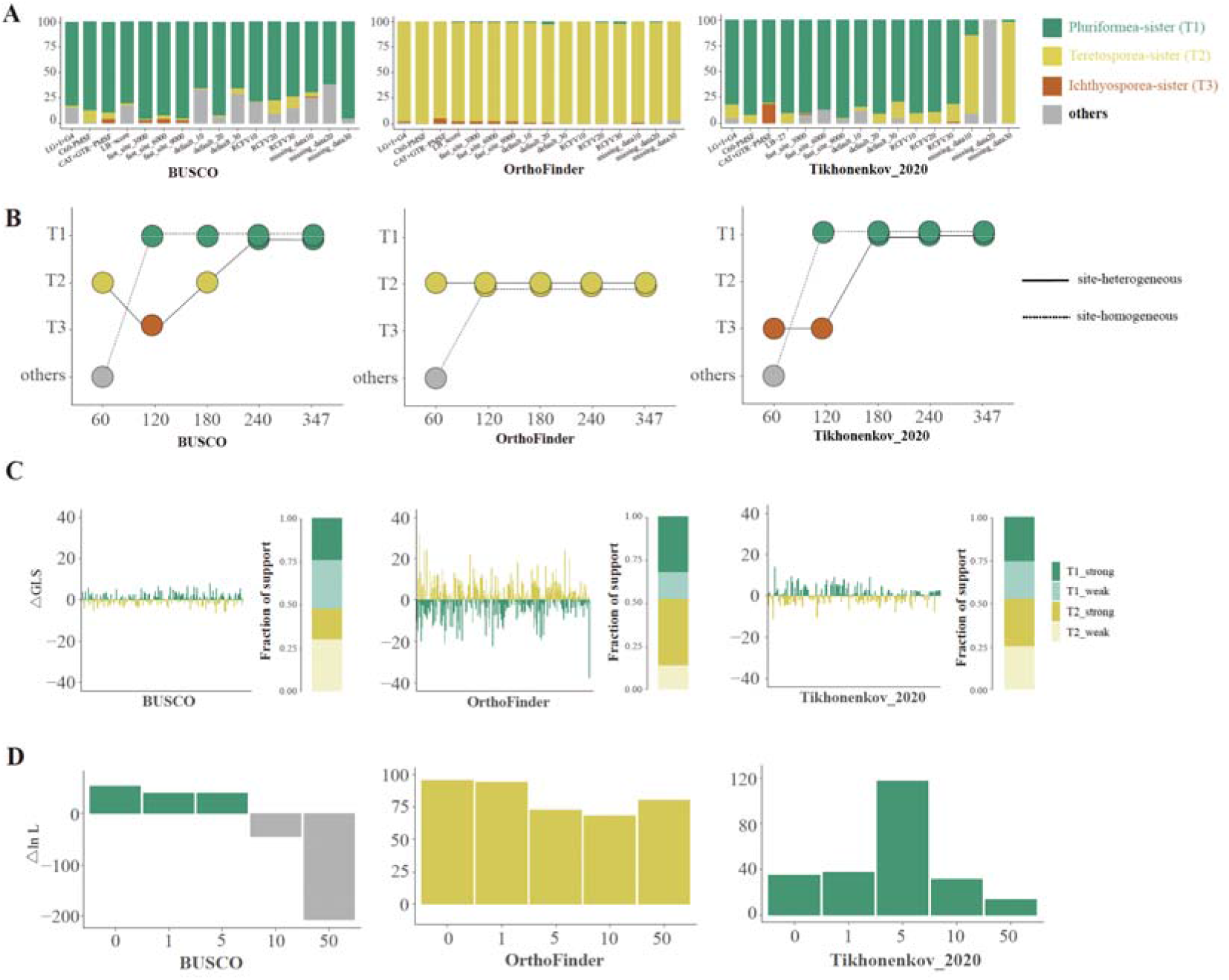
Multi-dimensional analysis of alternative phylogenetic hypotheses. **A**, Bootstrap support values for alternative hypotheses across different datasets are presented. The stack bar plots indicate the occurrence frequencies of each topology in 1000 UFB trees. **B,** Topological differences among different taxon- sampling densities and modeling schemes, also refer to Extended data Fig.4. **C,** Bar plot of the difference in gene log-likelihood scores (ΔGLS) between the two hypotheses recovered in this study. Proportions of genes supporting each of two alternative hypotheses for three data matrices are also shown. The ΔGLS values for the genes across each data matrix can be found in Supplementary Table 9. We considered a gene with an absolute value of log-likelihood difference of 2 as a gene with strong (|lnL| > 2) or weak (|lnL| < 2) phylogenetic signal. **D,** Quantification of the effect of the removal of tiny amounts of data on the branch of interest’s topology. For each branch, the 1, 5, 10, 50 genes with the highest absolute ΔGLS values were excluded; The y axis is the log-likelihood scores (Δln L) difference for the favored topological hypothesis. If Δ ln L > 0, then the resulting tree supports T1, if Δ ln L < 0, then the resulting tree supports T2 or ‘others’. Different hypotheses favored are marked by different colors, the green bars indicate the favored topology is T1, the yellow bars indicate the favored topology is T2, gray bars indicate the resulting trees supporting hypotheses other than T1, T2 and T3.

Previous studies of Opisthokonta phylogeny have emphasized that site-heterogeneous models are the most appropriate for deep phylogenomics^11,14,18,38,39^. These models incorporate a higher degree of biological realism by specifically accounting for compositional heterogeneity, a common feature among datasets spanning ancient evolutionary distances^68^. However, relationships among unicellular Holozoa were largely robust to modeling schemes suggesting compositional heterogeneity is not the dominant contributor to the incongruences observed. This aligns with the results from the subsampling analysis. The reduction in compositional heterogeneity within the data matrix, achieved by progressively removing genes with high RCFV scores, did not appear to influence the topologies (Figure 4A, Supplementary Table 5).

### Different gene sets are a source of incongruence in Holozoa

Sensitivity analysis—reinferring species-level relationships using 18 subsampling strategies—revealed high degrees of congruence within but not between data matrices (Extended data Fig. 3, Supplementary Figs. 19-57, Supplementary Table 6). Specifically, phylogenies inferred using subsets of the BUSCO, OrthoFinder, and Tikhonenkov_2020 shared 97.2%, 95.9%, and 97% of bipartitions, respectively, whereas the average bipartitions shared among submatrices from different data matrices were 88.0% (BUSCO versus Tikhonenkov_2020), 88.5% (OrthoFinder versus Tikhonenkov_2020) and 91.2% (BUSCO versus OrthoFinder) (Extended data Fig. 3, Supplementary Table 6).

This finding suggests different gene sets are the primary source of incongruence. Due to functional constraints and different evolutionary trajectories, genes may contain positions that undergo different degrees of multiple substitutions, resulting in varying saturation levels among datasets. The difference in saturation levels could contribute to the disparities in phylogenetic inference^47^. To test this hypothesis, we quantified the saturation level of the data matrices following Philippe et al.^47^ using PhyKIT^69^, data with no saturation will have a value of 1, while a value of 0 means complete data saturation. We found that the Tikhonenkov_2020 data matrices were the most saturated (∼0.12) and that the OrthoFinder data matrices were the least affected by multiple substitutions (∼0.24). The varying degrees of saturation may contribute to the observed distinctions among different data matrices (Supplementary Table 7). However, discerning the specific impact of saturation from other factors remains challenging.

Notably, the sensitivity analyses implicated missing data as a potential source of error for the Tikhonenkov_2020 matrices (Fig. 4A, Supplementary Table 5). Specifically, removing 10% and 30% of the genes with large amounts of missing data result in switch in support from Pluriformea-sister to Teretosporea-sister, however, remove 20% of the genes resulted in a topology likely to be erroneous (a clade composed of Filasterea, Ichthyosporea and Pluriformea), suggest the influence is not consistent and may due to the loss of signals. Identifying other predictors of topological preference is worthy of further investigation.

### Taxon sampling is vital for inferring Opisthokonta relationships

A prominent phylogenomic enigma is the placement of *T. unikontum*, the sole representative of a distinct Holozoa predator lineage with structural and molecular characteristics of evolutionary interest^18^. Our analyses mirrored this uncertainty (Supplementary Figs. 1-3, 7-9, 13-15). Exclusion of *T. unikontum* significantly improved nodal support of other unicellular holozoan lineages (Supplementary Figs. 4-6, 10-12, 16-18), suggesting *T. unikontum* is a rogue taxon. This finding was supported by graph-based analyses of the OrthoFinder matrix. Confident placement of rogue taxa can be achieved by sampling additional closely related taxa^49,70^, suggesting expanded taxon sampling may bring clarity to the placement of *T. unikontum*.

Examining the impact of taxon sampling revealed that insufficient taxon sampling led to uncertainties in the phylogenetic signal, resulting in erroneous or poorly supported inferences. For unicellular Holozoa, when down-sampling datasets to a number of taxa comparable to other studies^11,18,38,39^ (N_taxa_ = 60; data matrices #4), the phylogenetic inference using the LG model supported monophyly among Filasterea, Ichthyosporea, and Pluriformea, a topology that is widely refuted^14,39^ (Fig. 4B, Extended data Fig. 4A, Supplementary Figs. 58, 62, 66). Expanding taxon sampling to 120, 180, or 240 taxa (data matrices #4-7) yielded the same topology as the 347-taxon data matrices (Fig. 4B, Extended data Fig. 4A, Supplementary Figs. 59-61, 63-65, 67-69). Increased model complexity on sparsely taxon-sampled data sets introduced more uncertainty (Supplementary Figs. 70-81). For example, using the GTR+CAT model, the topology of Tikhonenkov_2020#4 switched to supporting the Ichthyosporea- sister hypothesis (Fig. 4B, Extended data Fig. 4A). Notably, this topology was also supported in a study using the same genes, similar taxon sampling, and highly complex substitution models. In contrast, using the same genes, similarly complex models, but improved taxon sampling (in this case, 180, 240, and 347 taxa), resulted in robust support for the Pluriformea-sister hypothesis irrespective of model complexity (Fig. 4B, Supplementary Figs. 18, 80-81). This observation suggests that the support for the Ichthyosporea-sister hypothesis may be an artifact of insufficient taxon sampling.

Additionally, our results show expanded taxon sampling improved the resolution and robustness of the phylogenetic inference. Specifically, we found that sparse sampled subsets failed to resolve some well-established relationships. For instance, OrthoFinder data matrices with 60 taxa (#4) and 120 taxa (#5) under both simple and complex models produced phylogenies inaccurately positioning Zoopagomycota as the sister group to Dikarya (Supplementary Figs. 58-59, 62- 63). Similarly, Tikhonenkov_2020 data matrices with 120 taxa (#5) and 180 taxa (#6), misplaced Xenacoelomorpha as more closely related to protostomes instead of as the sister clade to Bilateria (Supplementary Figs. 67-68, 79-80). Moreover, the average bootstrap value of trees based on submatrices showed an upward trend as the increasing number of taxa sampled. For example, the average bootstrap value based on BUSCO submatrices increased from 93.1 to 98.9 (Extended data Fig. 4B). Similar trends were observed for the OrthoFinder and Tikhonenkov_2020 submatrices.

The importance of taxon sampling for obtaining more accurate and reliable phylogenetic conclusions is widely accepted^71–76^. For example, a simulation study revealed a correlation between a decrease in taxon sampling and an increase in the average branch length of terminal branches, potentially exacerbating LBA artifacts^77^. The branches that separate non-animal Holozoa lineages are short and might therefore be especially sensitive to systematic errors. In cases where the number of sampled species is limited, the resultant non-phylogenetic signal can overwhelm the faint genuine phylogenetic signal present in the ancient divergence of the Holozoa phylogeny. In line with this, the results of Tikhonenkov et al. 2020 noted that taxon sampling could be a significant factor regarding the resolution of unicellular Holozoa; perturbing taxon sampling in the same data matrix changed inferred topologies. Nonetheless, our analyses, together with previous reports^48–50,78^, suggest expanded taxon sampling is a promising approach to address contentious evolutionary relationships among Opisthokonta. Future sampling strategies may focus on under-sampled lineages and “breaking” long branches^79^.

### Distributions of single-gene support highlight the importance of genome-scale approaches

The distribution of concordance factors (measures for quantifying genealogical concordance in phylogenomic datasets) across branches of two contrasting topologies further proved that the relationships of unicellular Holozoa are among the most difficult to resolve. Specifically, contentious nodes consistently had low gCF scores but high UFB support (Extended data Fig. 5, Supplementary Table 8). For instance, despite the Teretosporea-sister hypothesis being strongly supported using OrthoFinder#2 matrix under a site-homogeneous model (UFB support = 98), gCFs revealed only 0.7% (3 / 426) of individual loci supported the Teretosporea-sister hypothesis. Furthermore, 98.6% (420 / 426) of gene trees supported topologies other than T1, T2, and T3, revealing low concordance among gene trees. Examining sCF revealed substantial noise among single sites evidenced by a similar proportion of support for each hypothesis (34.04 / 32.98 / 32.98; Supplementary Table 8). Consequently, the incongruences are likely due to weak phylogenetic signals in single loci.

Distributions of phylogenetic signals, which use gene likelihood scores (ΔGLS values) to compare support for two topologies, also revealed weak single-gene phylogenetic signals (Fig. 4C, Supplementary Table 9). Specifically, across the three data matrices, the proportions of genes supporting Pluriformea-sister (T1) or Teretosporea-sister (T2) were similar, albeit contrasting—in the BUSCO#2 matrix, 52.6%(120 / 228 genes) supported T1, and 47.4% (108 / 228 genes) supported T2; in the OrthoFinder #2 matrix, 47.7% (210 / 440 genes) supported T1, and 52.3% (230 / 440 genes) supported T2; and in the Tikhonenkov_2020#2 matrix 52.2% (105 / 201 genes) supported T1, and 47.8% (96 / 201 genes) supported T2 (Fig. 4C, Supplementary Table 9).

Inferences were largely insensitive to removing the small amount of data. Pruning the 1, 5, 10, and 50 genes with the highest absolute ΔGLS values revealed that the OrthoFinder#2 and Tikhonenkov_2020#2 matrices successfully recapitulated evolutionary relationships inferred using the full data matrix, indicating support for the two topologies was derived from many genes. In contrast, the Pluriformea-sister hypothesis was not supported when 10 or more genes in the BUSCO#2 matrix were removed (Fig. 4D, Supplementary Table 10), suggesting a small subset of orthologs may have a disproportionate influence on phylogenomic inference warranting cautious interpretation of results. Single-gene sensitivity may be attributed to the BUSCO#2 matrix harboring the weakest phylogenetic signal (average |ΔGLS|=1.91), as compared the OrthoFinder#2 and Tikhonenkov_2020#2 matrices (average |ΔGLS| = 5.33 and 2.68, respectively; Fig. 4C, Supplementary Table 9).

Taken together, examining distributions of single-gene support reveals weak signals in single loci and sites therein, underscoring the importance of genome-scale approach to tree of life inquiries. Future studies will likely benefit from expanded taxon and gene sampling to address uncertain relationships among unicellular Holozoa, especially among key lineages and close relatives.

## Conclusions

Recent comparative genomics studies suggest that the unicellular Holozoa already had a rich repertoire of genes required for cell adhesion, cell signaling, and transcriptional regulation in modern animals^2,11,39,67^. A comparative study that supported the Ichthyosporea-sister hypothesis suggests an animal-like extracellular matrix emerged after Ichthyosporea diverged from all other Holozoa, suggesting a single origin^11^. However, if Pluriformea were to be considered the sister lineage to the rest of the Holozoa, it would suggest secondary losses of the animal-like extracellular matrix in the unicellular lineages. Thus, the branching order of unicellular relatives of animals is critical for interpreting the sequence of events that led to the emergence of animals and their potential contributions to the origin of multicellularity. Nevertheless, the precise evolutionary origins of these traits remain an area of ongoing exploration.

Resolving rapid divergences in deep time can be extremely difficult^80–82^. Branches that capture divergences among unicellular Holozoa are short and buried deep in the Opisthokonta tree of life. Sources of error resulting in low signal-to-noise ratios, like saturation by multiple substitutions, can be masked by undue confidence in topologies inherent to the concatenation approach. Despite these challenges, we devised a phylogenomic approach (Fig. 2C) able to uncover artifacts contributing to errors and evaluate the robustness of phylogenomic inference. We found that denser taxon sampling can significantly improve the robustness of the Opisthokonta tree of life, and previous support for the Ichthyosporea-sister hypothesis likely stems from insufficient taxon sampling (Fig. 4B, Extended data Fig. 5). In contrast, we show that different gene sets are the main contributor to incongruence between the Pluriformea-sister and Teretosporea-sister hypotheses (Extended data Fig. 3). Thus, expanded taxon and gene sampling are promising solutions to illuminate early Holozoa evolution. Incorporating other sources of phylogenomic information, such as rare genomic changes^83^ like macrosynteny^26,84,85^, may also prove insightful.

In summary, we inferred a well-supported Opisthokonta Tree of Life in geologic time (Fig. 3A), identified branches warranting further investigation, and uncovered promising strategies for resolving incongruence. Together with other studies, three topologies have received support concerning the root of the Holozoa Tree of Life (Fig. 2B), suggesting this may be one of the most challenging enigmas in the phylogenomic era. We anticipate our results will be useful in studying genes and phenotypes across Opisthokonta in geologic time. Moreover, methods developed herein may facilitate careful investigations into the Tree of Life.

## Methods

### Data acquisition

Genome and transcriptome data for over eight hundred Opisthokonta species were retrieved from public databases. Transcriptome data were included due to the limited availability of genomic data for certain lineages, such as unicellular holozoans, Ctenophora, Porifera, and Cnidaria. Representatives of fast-evolving lineages containing pathogens and parasites known to cause long-branch attraction (LBA) were excluded (i.e., Microsporidia, Platyhelminthes, Nematoda)^27,86^.

To minimize the amount of missing data and remove potential low-quality genomes/transcriptomes, completeness was assessed using the Benchmarking Universal Single-Copy Orthologs (BUSCO) v5.02^87^ pipeline with the eukaryotic_odb10 database (255 near-universally single-copy orthologs or BUSCO genes; last accession date: June 14, 2022)^88^. BUSCO genes were classified as single-copy, duplicated, fragmented, or missing based on the presence/absence, copy number, and length of the predicted BUSCO gene; the fraction of single-copy BUSCO genes present is a proxy for assembly completeness. With the exception of unicellular lineages and non-bilaterian animal lineages, other taxa were filtered based on BUSCO gene completeness while also ensuring a balanced representation of different Opisthokonta lineages. The final list contained 339 Opisthokonta species (217 genomes and 122 transcriptomes). Additionally, nine outgroup taxa were downloaded from NCBI (Last accession date: December 17, 2022) based on the current understanding of Opisthokonta phylogeny^14,38^ (Supplementary Table 1).

### Construction of three phylogenomic data matrices

To explore the effectiveness of different ortholog inference methods, we constructed two novel data matrices using different strategies—that is, targeted identification of phylogenomic markers (BUSCO) and de novo inference (OrthoFinder)—and an additional one that built on a data matrix employed in an early phylogenomic analysis^18^ (Fig. 2C).

#### (i) BUSCO data matrix

BUSCO genes have proven to be useful phylogenomic markers in diverse lineages^43,87,89^. Therefore, a data matrix was constructed using complete, single-copy sequences identified with the BUSCO algorithm as described above, resulting in 255 single-copy orthologs.

#### (ii) OrthoFinder data matrix

Orthologous groups were initially constructed using the genomic data from 52 taxa—49 Opisthokonta species and three outgroup taxa. Each major Opisthokonta lineage was represented by one to three taxa with the best assembly quality (Supplementary Table 11). OrthoFinder v2.5.4 ^90^ was used to identify putatively orthologous sequences shared among taxa using default parameters (inflation parameter 1.5). To identify additional phylogenomic makers, species-specific inparalogs—genes that have undergone duplication events along terminal taxa—were pruned from groups of orthologous genes^91,92^. To do so, orthogroups with greater than or equal to 80% taxon occupancy (N = 42) were aligned with MAFFT v7.505^93^ using the auto parameter and maxiterate set to 1000. Ambiguously aligned sites were removed using trimAl v1.415^94^ with the “gappyout” option following benchmarking studies^95,96^. Approximate Maximum Likelihood (ML) phylogenies were inferred from the trimmed alignments using FastTree v2.2.11 with the slow and gamma arguments^97^. Species-specific inparalogs were trimmed using PhyloPyPruner v0.9.5 (https://pypi.org/project/phylopypruner) with the following arguments: “--min-len 50 --trim-lb 7 --min-support 0.75 --min-taxa 35 --trim-freq-paralogs 5 --prune LS”, resulting in 635 single-copy orthologs. A profile Hidden Markov Model (HMM) was made for each single-copy ortholog using hmmbuild in HMMER v3.2.1^98^.

The resulting HMMs and orthofisher v1.0.3^99^ were used to identify single-copy orthologs in the 348 proteomes using a fractional bitscore threshold of 0.95.

#### (iii) Tikhonenkov_2020 data matrix

Another data matrix was constructed using a set of 201 previously identified Opisthokonta orthologs^18^. As described above, HMMs were built from the multiple sequence alignments using HMMER, and orthofisher was used to identify single-copy orthologs in each proteome.

#### Supermatrix construction

Single-copy orthologs from each data set were treated using the same procedure (Fig. 2C). Specifically, quality filtering for unaligned single-copy ortholog sequences was done using PREQUAL v1.02^100^ with a 0.95 posterior probability filtering threshold. Filtered sequences were then aligned with MAFFT v7.505 ^93^ using the argument globalpair, maxiterate set to 1000, and unalignlevel set to 0.6. Alignments were then processed with Divvier v1.01^101^ using the “divvygap” option and requiring a minimum of four characters per column for output. Multiple sequence alignments with lengths less than half of the total alignment length were removed. Highly divergent and gappy sites (>80% gaps) were then trimmed using BMGE v.1.12.2 with default settings^102^. Multiple sequence alignments shorter than 100bp or with less than 70% taxon representation were removed. Remaining multiple sequence alignments were concatenated using PhyKIT v1.11.10^69^. The final BUSCO, OrthoFinder, and Tikhonenkov_2020 data matrices contained 228, 440, and 201 genes, respectively, and are represented as BUSCO#1, OrthoFinder#1, and Tikhonenkov_2020#1 (Fig. 2C, Supplementary Table 2). The overlap between the three data matrices was identified using an all-versus-all comparison using DIAMOND^103^. Functional categories of each ortholog set in three data matrices were annotated using eggNOG v5.0^104^ and BLASTP searches.

### Phylogenomic analysis

To infer the Opisthokonta phylogeny and evaluate the impact of different models on the tree topology, we performed phylogenetic analyses using both site-homogeneous and site-heterogeneous evolutionary models (Fig. 2C). The best-fitting substitution model (LG) was determined using ModelFinder^105^ with the option msub set to nuclear. We first inferred phylogenetic trees using the computationally efficient site-homogeneous model LG+I+G4 (hereafter referred to as LG). For site-heterogeneous models, the large size of our data matrices is intractable for the C models^106^ and the CAT model^107^ implemented in IQ-TREE and PhyloBayes, respectively. However, approximations thereof offer similar benefits and require fewer, but still substantial, resources. Thus, we employed the PMSF (posterior mean site frequency) approximation for these two models, which requires a guide tree (inferred using the site-homogenous mode), site-specific stationary distributions, and amino acid exchangeabilities. Approximate site-specific stationary distributions and amino acid exchangeabilities were estimated using the Bayesian GTR+CAT-PMSF model^107,108^ (hereafter refer to as GTR+CAT) with 1,100 generations and a burn-in of 100 using PhyloBayes-MPI^109^ following a previous study^110^. Results were reformatted using publicly available scripts (https://github.com/drenal/cat-pmsf-paper) to be compatible with IQ-TREE 2. Tree inference was then performed in IQ-TREE 2 using the LG+C60+F+G4 model under the PMSF approximation (hereafter referred to as LG+C60)^106,111^.

For each dataset, branch support was evaluated using ultrafast bootstrap (UFB) replicates. Using 1000 UFB replicates^112^, branch support was binned into three categories: strongly supported (above 95), moderately supported (between 90 and 95), and weakly supported (below 90) following a previous study^113^.

### Molecular dating

To infer the timing of Opisthokonta divergences, we used the Bayesian method MCMCTree in the paml4.9e package^114^. MCMCTree analyses were run on the OrthoFinder#1 data matrix using approximate likelihood calculations with (i) uncorrelated (clock = 2) and (ii) autocorrelated (clock = 3) relaxed clock models and the topology inferred using the LG+C60 model. We used ten node calibrations based on well-established fossil evidence—seven from Metazoa and three from fungi^60,115–119^ (Supplementary Table 4). For computational tractability, MCMCTree were run on 10 replicate data matrices, each consisting of a randomly chosen subset of 100 genes. Two independent MCMC chains were run for the analysis, each consisting of 1.5 million generations, discarding the first 100,000 generations as burn-in. Lastly, the divergence time estimate for each internal branch was calculated as the average across the timetrees produced by the 10 replicates.

### Systematically Evaluating Analytical Errors

Phylogenetic inference of deep divergences, such as those concerning major Opisthokonta lineages, are susceptible to many sources of error that may lead to erroneous reconstructions^70,80,86,120,121^. By combining information theory, which enables quantifying the phylogenetic informativeness of orthologs, and sensitivity analyses, we conducted a systematic evaluation of sources of error^19,25,29,110^. Specifically, a series of submatrices were generated using an information theory-based framework. Subsetting strategies featured subsampling taxa, sites, or genes based on multiple dimensions of information content, such as rogue taxa, long branch scores (LBS), rates of sequence evolution, composition heterogeneity (measured by relative composition frequency variability or RCFV;^122,123^, missing data, and phylogenetic usefulness, a metric that incorporates multiple dimensions of information^110^ (Fig. 2C). We also tested the effect of taxon sampling on the resolution of unicellular Holozoa using a taxonomy-informed subsampling strategy (Fig. 2C). The details of data matrices generated in the analyses can be found in Supplementary Table 2. To remove potential confounding effects, subsetting was conducted on the rogue taxon pruned data matrices (denoted by the suffix “#2”, see below).

#### Rogue Taxa – data matrices #2

Rogue taxa are known to cause phylogenomic incongruence^70^. Rogue taxa were identified in the three full data matrices (denoted by the suffix “#1”) using RogueNaRoK^124^, revealing *T. unikontum* is a putatively rogue taxon in the OrthoFinder#1 data matrix, but not the other two data matrices. This observation corroborates previous reports that the placement of *T. unikontum* is unstable and its inclusion leads to unresolved trees^18^. To examine the effect of rogue taxon exclusion, *T. unikontum* was pruned from each data matrix (Supplementary Table 2). We then performed the same phylogenetic analyses as described above on the resulting data matrices.

#### Long branch score – data matrices #3

LBS, a metric that can be used to identify taxa that might cause LBA^125^), was calculated for each taxon using PhyKIT^69^ following^125^. This analysis identified 27 “long-branched” taxa (Supplementary Table 12), which were pruned from the #2 data matrices (Fig. 2C).

#### Taxon sampling - data matrices #4-7

To assess the impact of taxon sampling on phylogenetic topologies, four submatrices with different taxon sampling densities (while maintaining a high diversity) were generated. To be comparable with the taxon number in previous studies (Torruella et al. 2015, 83 taxa; Grau-Bove et al. 2017, 57 taxa; López-Escardó et al. 2019, 79 taxa; Hehenberger et al. 2017, 38 taxa; Tikhonenkov et al. 2020, 75 taxa), 60 taxa representing 25 major lineages in Opisthokonta were selected while preserving the most comprehensive representation of Filasterea, Ichthyosporea, and Pluriformea (Supplementary Table 13). The impact of increased taxon sampling was evaluated by randomly selecting additional, non-redundant species from the remaining taxa to create three additional datasets of 120, 180, and 240 taxa (Supplementary Table 13), resulting in 12 new data matrices (Fig. 2C).

#### Fast evolving sites – data matrices #8-10

Fast-evolving sites may suffer from saturation by multiple substitutions and cause LBA artifacts^11^. For each data matrix, 3000, 6000, or 9000 sites with the highest rates of sequence evolution were removed using the fast_site_remover.py script from PhyloFisher^126^, which uses DistEst^127^ to estimate evolutionary rates, resulting in a total of nine new data matrices (Fig. 2C).

#### Phylogenetic usefulness – data matrices #11-13

Gene properties related to potential phylogenetic usefulness and bias were calculated using the genesortR package^110^. Phylogenetic usefulness was determined using principal component analysis of seven gene properties: Robinson-Foulds distance to the best estimated species tree; average bootstrap support; saturation; compositional heterogeneity; root to tip variance; average patristic distance; and proportion of variable sites. The three data matrices were then subsampled using the best scoring 90, 80, and 70 percent of genes (Fig. 2C).

#### Compositional heterogeneity – data matrices #14-16

Compositional heterogeneity has been implicated as an important source of systematic error in Opisthokonta phylogeny^14,18,65,128^. The 90, 80, and 70 percent of genes with the lowest RCFV scores, indicative of being least prone to compositional biases, were subsampled using genesortR (Fig. 2C).

#### Missing data – data matrices #17-19

To examine the impact of missing data (that is, taxon occupancy), the 90, 80, and 70 percent of genes with the least missing data were subsampled using genesortR (Fig. 2C).

#### Phylogenetic inference of subsetted data matrices #3-19

We performed ML phylogenetic analyses with IQ-TREE 2^129^ on the subsampled matrices using a single LG model, assessing topological support with 1000 UFBs^112^. Phylogenetic inference of data matrices #4-7 were further examined using the GTR+CAT model as described above. Support for the three alternative topologies (Pluriformea-sister, Teretosporea-sister and Ichthyosporea-sister hypotheses, correspond to T1, T2 and T3, respectively, Fig. 2B) was also examined by examining the frequency of each topology among the 1000 UFB replicates using IQ-TREE 2. Specifically, cladogram of T1: (Pluriformea, (Ichthyosporea, Filozoa)), T2: ((Pluriformea, Ichthyosporea), Filozoa) and T3: (Ichthyosporea, (Pluriformea, Filozoa)) were input to IQ-TREE 2 via the sup option, with the remaining taxa constrained as polytomies. Over 214,489 central processing unit hours (∼24.5 years) were used for this study.

### Quantifying single-gene phylogenetic signal

Single-gene phylogenetic signal was quantified using two approaches: likelihood scores and concordance factors. Gene concordance factors (gCFs) and site concordance factors (sCFs)—the percentage of gene trees that support a node based on descendant taxa and the percentage of informative sites that support that node via parsimony, respectively—were calculated using IQ-TREE 2. To calculate gCFs, individual gene trees were first inferred using IQ-TREE 2 using the best fitting substitution model selected by ModelFinder with the msub parameter set to nuclear, gCFs were then estimated by comparing individual gene trees to the concatenated tree inferred with LG model; sCFs were calculated using 100 random quartets.

To examine phylogenetic signals supporting two conflicting hypotheses recovered in this study (T1 and T2, see Fig. 2B), we examined the gene likelihood scores for each data matrix (#2). Site-wise support was calculated for both hypotheses using IQ-TREE 2 with the g option and the LG model. The numbers of genes supporting each hypothesis were then calculated from IQ-TREE 2 using the wsl option by comparing genewise log-likelihood scores (ΔGLS)^130^. Genes with an absolute value of log-likelihood difference greater than two (|ΔlnL| > 2) were considered to have strong phylogenetic signal; those with a difference less than two (|ΔlnL| < 2) were considered to have weak signals.

To examine the influence of single genes with high ΔGLS values, each of the data matrices #2 were subsampling by pruning the 1, 5, 10, and 50 genes with the highest absolute ΔGLS values following Shen et al.^130^, resulting in 12 new data matrices. A species tree was then estimated for each matrix using IQ-TREE 2 with the LG model and 1000 ultrafast bootstrapping replicates^112^.

---

**Extended data Fig. 1:**
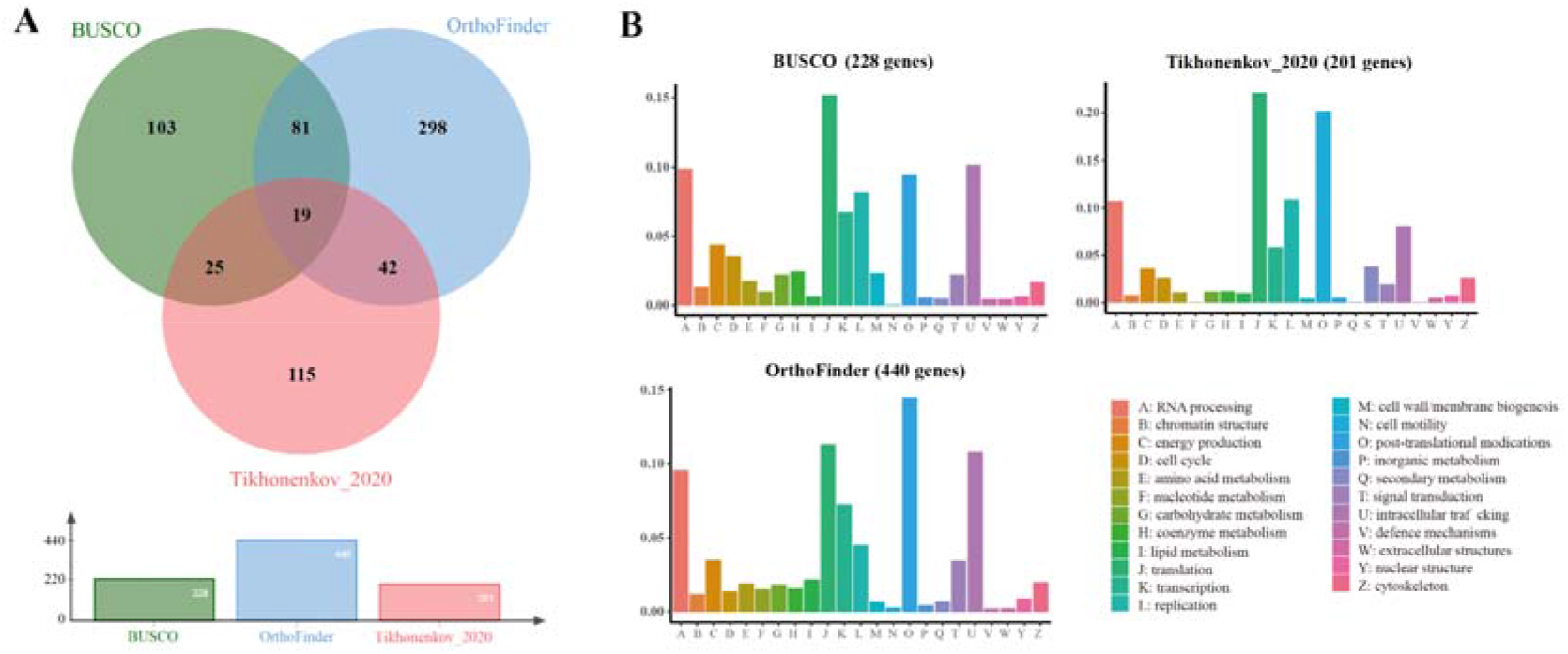
Comparison of the three data matrices constructed in this study. **A**, Venn diagram of shared orthologs for the three data matrices. The BUSCO data matrix overlaps with the OrthoFinder data matrix by 100 genes, and by 44 genes with the Tikhonenkov_2020 data matrix, see also Supplementary Table 15. **B**, Single copy orthologs with functional information, the functional category ‘S: unknown function’ was ignored as it does not include functional information. The functional categories of every gene were determined by averaging the annotations of the corresponding cluster members.

**Extended data Fig. 2:**
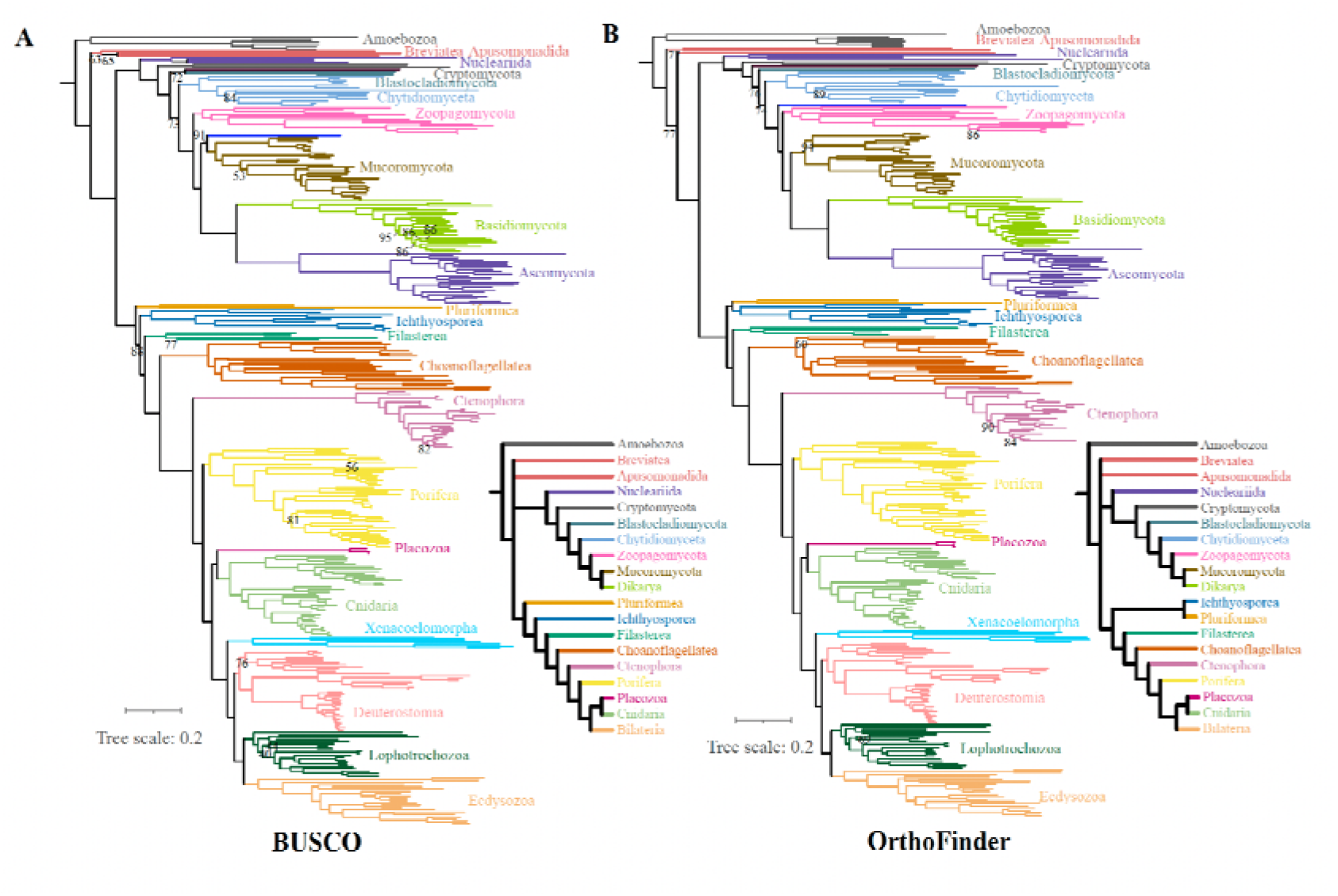
Comparison of the phylogenetic inference using BUSCO and OrthoFinder data matrices under site heterogeneous model. **A**, The topology of the IQ-TREE 2 inference with the BUSCO data matrix#2 using the LG+PMSF(C60)+F+G4 model. **B**, The topology of the IQ-TREE 2 inference with the OrthoFinder data matrix#2 using the LG+PMSF(C60)+F+G4 model.

**Extended data Fig. 3:**
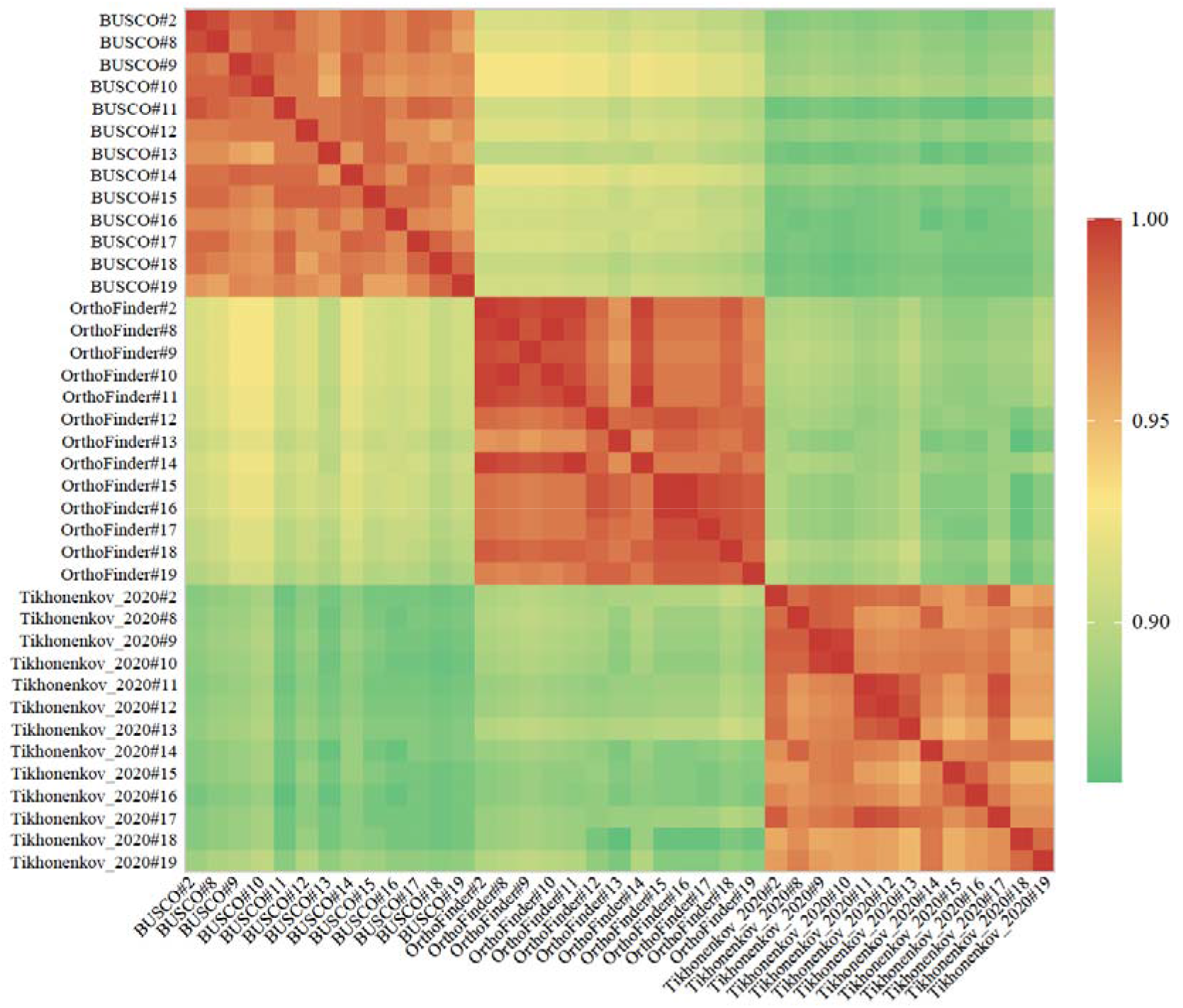
Heatmap of topological similarities for all pairwise comparisons among the phylogenies reconstructed from subsampling analyses of 39 data matrices (#2, #8-19) during sensitivity analysis. The topological congruence between each pair of phylogenies was calculated using gotree^131^, function “compare”. The color of the squares represents the number of bipartitions shared between trees. Results from data matrices#3-7 are not shown here since they do not share the same number of tree tips.

**Extended data Fig. 4:**
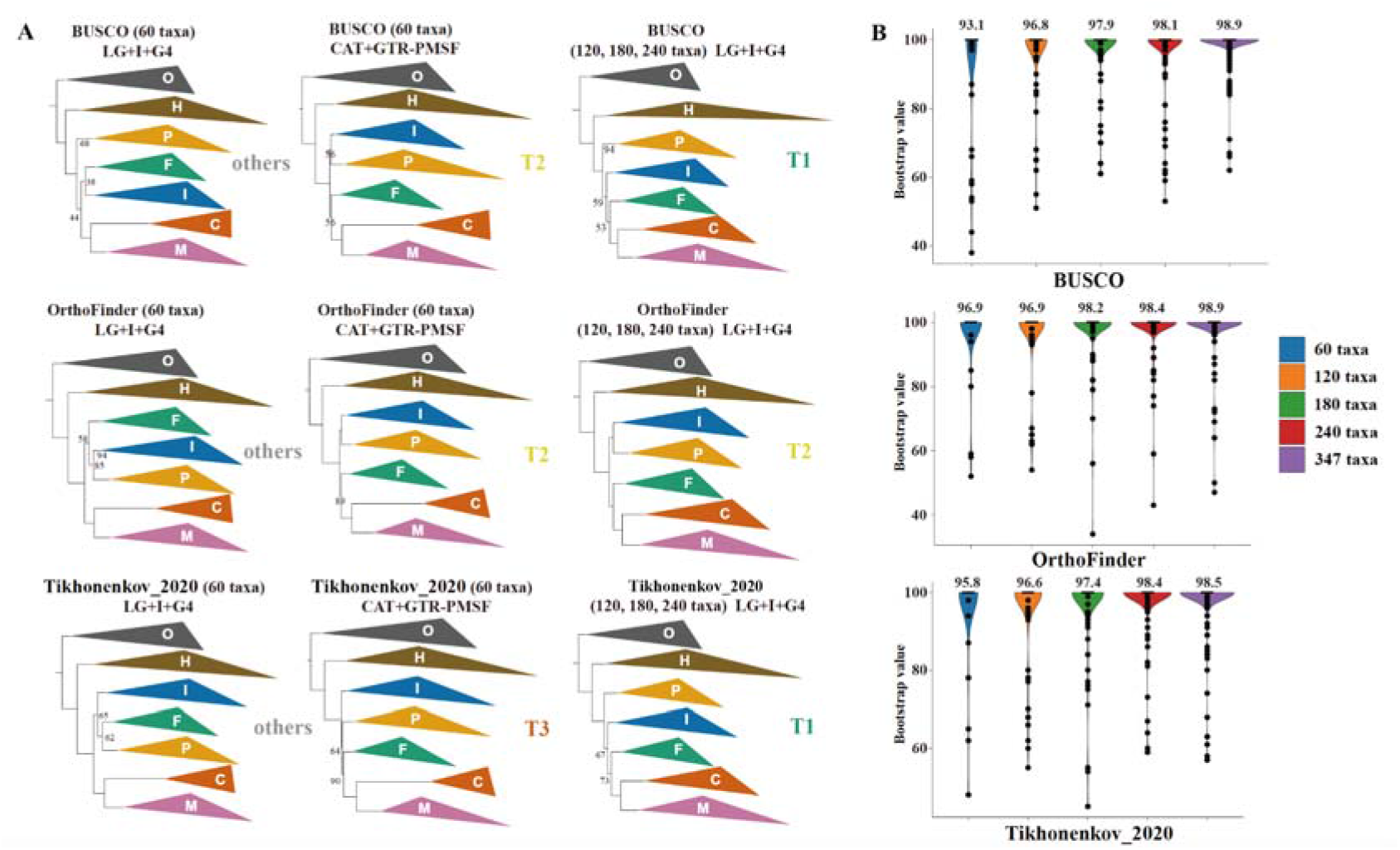
Comparison of topological differences among different taxon sampling densities. **A,** ML inference of data matrices with different sampling density, models used were labeled correspondingly. Lineages are abbreviated to the first letter of the taxon name, while “O” stands for the outgroup taxa. “M’’ refers to Metazoa, “C” refers to Choanoflagellatea, “F” refers to Filasterea, “I” refers to Ichthyosporea, “P” refers to Pluriformea, “H” refers to Holomycota. The bootstrap values of ML inference with data matrices#5-7 using the LG model are from corresponding data matrix #7, Nodes with bootstrap support >95 are not shown. **B**, Distribution of ultrafast bootstrap values in data matrices#2, 4-7, with average bootstrap values labeled.

**Extended data Fig. 5:**
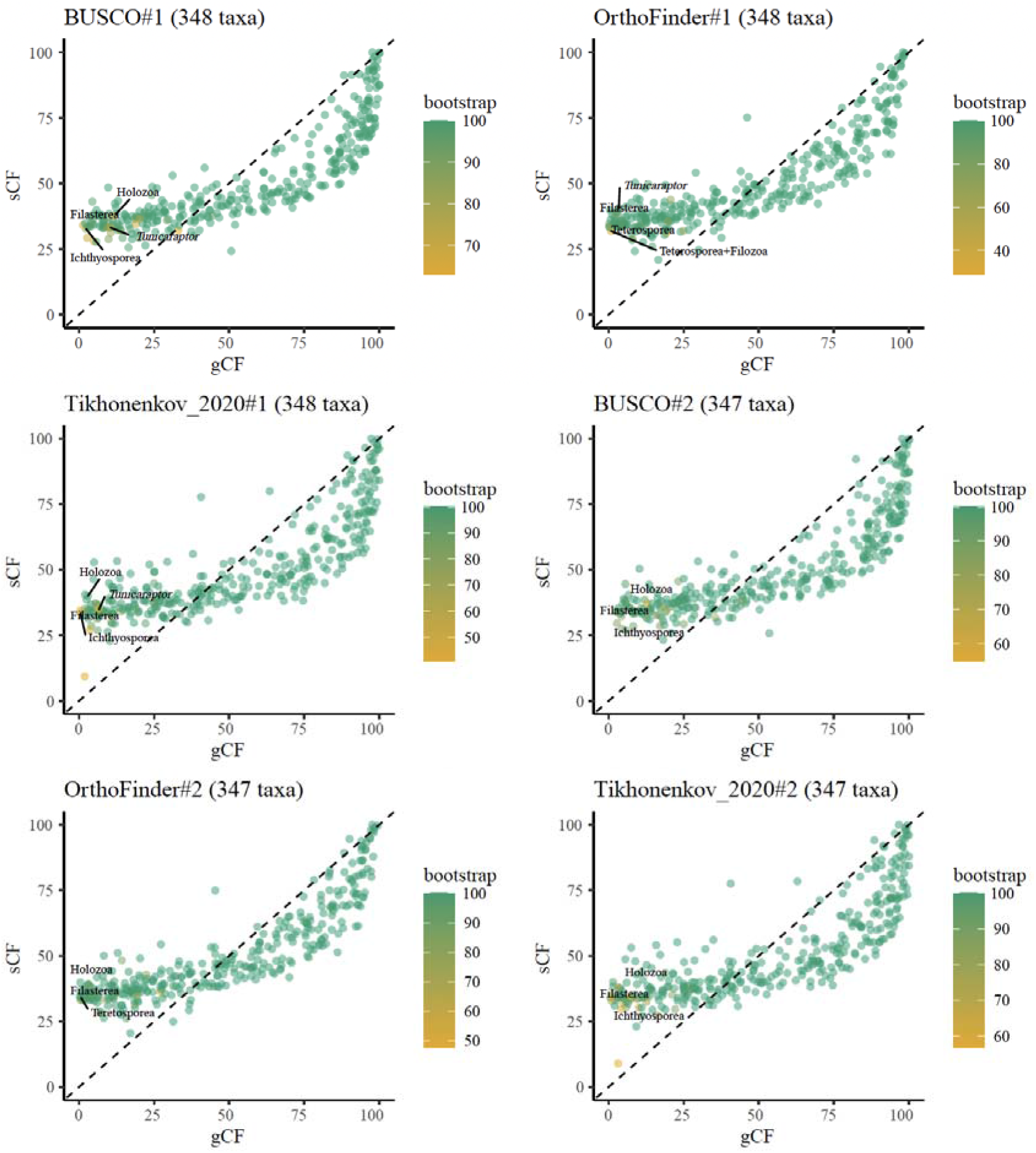
The distribution of gene concordance factors(gCFs) and site concordance factors(sCFs) across all nodes of the tree. Critical nodes concerning the relationships of unicellular Holozoa were labeled. The script for this plot is available at http://www.robertlanfear.com/blog/files/archive-2018.html.

